# Measuring renal cortical cell-specific mitochondrial metabolism

**DOI:** 10.1101/2024.11.24.622516

**Authors:** Kyle Feola, Andrea H. Venable, Mina Rasouli, Julie Do, Tatyana McCoy, Claire B. Llamas, Dana Straus, Prashant Mishra, Behzad Najafian, Sarah C. Huen

## Abstract

The metabolic health of the kidney is a primary determinant of the risk of progressive kidney disease. Our understanding of the metabolic processes that fuel kidney functions is limited by the kidney’s structural and functional heterogeneity. As the kidney contains many different cell types, we sought to determine the intra-renal mitochondrial heterogeneity that contributes to cell-specific metabolism. To interrogate this, we utilized a recently developed mitochondrial tagging technique to isolate kidney cell-type specific mitochondria. Here, we investigate mitochondrial functional capacities and the metabolomes of the early and late proximal tubule (PT) and the distal convoluted tubule (DCT). The conditional MITO-Tag transgene was combined with *Slc34a1-CreERT2*, *Ggt1-Cre*, or *Pvalb-Cre* transgenes to generate mouse models capable of cell-specific isolation of hemagglutinin (HA)-tagged mitochondria from the early PT, late PT, or the DCT, respectively. Functional assays measuring mitochondrial respiratory and fatty acid oxidation (FAO) capacities and metabolomics were performed on anti-HA immunoprecipitated mitochondria from kidneys of ad libitum fed and 24-hour fasted male mice. The renal MITO-Tag models targeting the early PT, late PT, and DCT revealed differential mitochondrial respiratory and FAO capacities which dynamically changed during fasting conditions. The renal MITO-Tag model captured differential mitochondrial metabolism and functional capacities across the early PT, late PT, and DCT at baseline and in response to fasting.

**New & Noteworthy:** This study described the generation and utilization of mouse models capable of interrogating kidney tubular epithelial cell-specific mitochondrial metabolism. Applying the MITO-Tag system in the kidney, we have for the first time defined the mitochondrial metabolic heterogeneity of renal cortical tubular epithelium and discovered differential mitochondrial functional capacities in response to an acute metabolic stress such as fasting.

## Introduction

Mitochondrial dysfunction has been implicated as a significant driver of disease pathology, including acute and chronic kidney disease. Recent studies have interrogated the differences of mitochondria across tissues and found significant heterogeneity in protein composition and substrate utilization, suggesting tissue-specific mitochondrial characteristics (1, 2). As the kidney’s cellular composition is diverse, it is likely that intra-renal mitochondrial heterogeneity contributes to kidney cell-specific mitochondrial function. Thus, an investigation of kidney cell-specific mitochondrial heterogeneity is needed to fully understand the mechanisms by which mitochondrial dysregulation promotes kidney disease.

The kidneys are extremely metabolically active (3), having the second highest mitochondrial density compared to other tissues (4). While comprising 1% of total body weight, the kidneys receive 25% of cardiac output, filter 180 liters of blood daily, and consume 10% of the total body oxygen for solute transport (to maintain electrolyte, pH, and volume homeostasis) and biosynthesis (hormone production for blood pressure, red blood cell, and bone homeostasis). To perform these functions, the kidney has at least 30 distinct cell types (5). The nephron, the basic unit of the kidney, is comprised of multiple tubular segments, the organization of which are critical to their diverse physiologic functions. Within the kidney, mitochondrial density (6) and function varies across the nephron (7). Metabolic studies often use cortical slices to interrogate renal cortical metabolism. However, it is likely that cortical tubular cells, including the proximal tubule (PT) and the distal convoluted tubule (DCT), are metabolically unique as they are functionally distinct (**Figure 1A,B**). The PT is further differentiated into separate segments (S1-S3), with the early segments responsible for initial bulk transport and the S3 segment most vulnerable to ischemic and nephrotoxic insults. Similarly, the DCT is differentiated into the early and late DCT (8). Due to nephron structure, parts of the PT and the DCT are physical neighbors within the renal cortex. The metabolic profiles of these distinct cortical tubular epithelial cells are therefore difficult to investigate in a cell-specific fashion. Current methods of interrogating kidney mitochondrial function include whole tissue mitochondrial isolation, microdissection of tubular segments, and single cell isolation using flow cytometric cell sorting with improved targeting of specific cell populations. While whole kidney mitochondrial isolation does not allow for cell-type specificity, micro dissection and flow cytometric cell sorting are limited by the need for tissue digestion and introduction of oxidative stress during the sorting process (9). We thus sought to use the MITO-Tag system in the kidney to investigate mitochondrial properties in a kidney cell-specific manner and to determine whether mitochondrial function differed among the early PT, late PT and the DCT.

**Figure 1.**
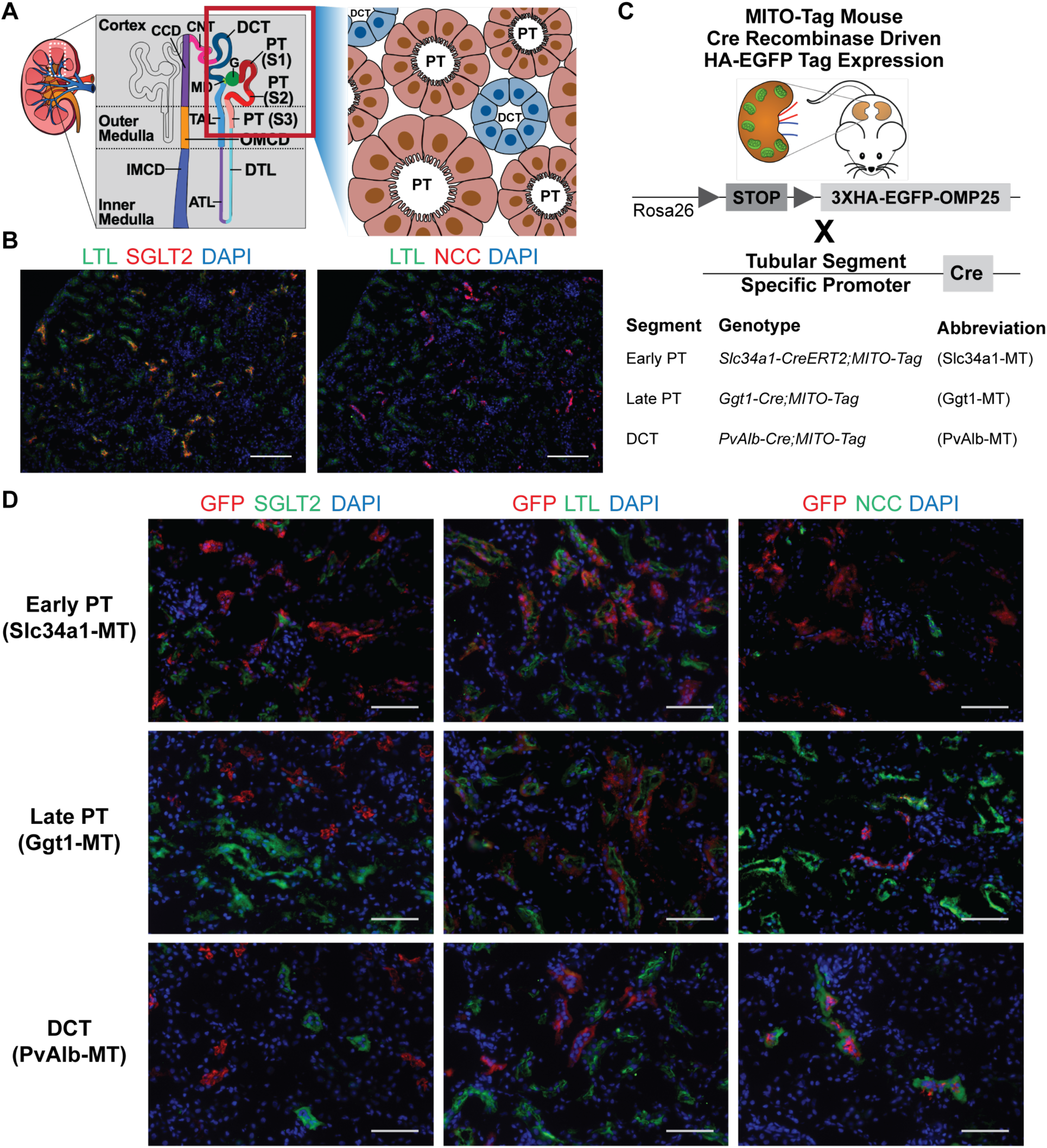
A model to study renal tubular mitochondrial heterogeneity. **A.** Schematic of the heterogeneous renal tubular epithelium. Inset highlights the juxtaposition of proximal tubular (PT) and distal convoluted tubular (DCT) cells. **B.** Immunostaining with sodium glucose co-transporter 2 (SGLT2, early PT S1-S2 segment), Lotus lectin (LTL, PT S1-S2>S3) and sodium chloride co-transporter (NCC, DCT). Scale bar = 100 μm. **C.** MITO-Tag mouse models to target the Early PT, Late PT, and DCT. **D.** Immunostaining with anti-GFP (anti-HA MITO-Tag includes GFP construct), SGLT2, LTL, and NCC of Slc34a1-MT, Ggt1-MT, and PvAlb-MT kidneys. Scale bar = 50 μm.

## Materials and Methods

### Mouse Studies

All animal experiments were performed in accordance with institutional regulations after protocol review and approval by the Institutional Animal Care and Use Committee of University of Texas Southwestern Medical Center. The MITO-Tag, (B6.Cg-Gt(ROSA)26Sortm1(CAG-EGFP)Brsy/J, mice (10) were bred with *Slc34a1-CreERT2* (11), *Ggt1-Cre* (12), or *Pvalb-Cre* (13) mouse lines to generate mouse models with cell type specific mitochondrial tags. MITO-Tag (RRID:IMSR_JAX:032290), *Ggt1-Cre* (RRID:IMSR_JAX:012841), and *Pvalb-Cre* (RRID:IMSR_JAX:017320) mice were purchased from Jackson Laboratory. *Slc34a1-CreERT2* (RRID:IMSR_JAX:032285) was obtained through the UT Southwestern O’Brien Research Core Center. *Slc34a1-CreERT2;MitoTag^+/-^*mice were gavaged with a single dose of tamoxifen in peanut oil (66 mg/kg body weight) and allowed a week to rest before experimental use. All mouse strains were maintained on a C57BL/6 background and housed under standard laboratory conditions with a 12:12-h light-dark cycle. Male mice 12 to 16 weeks old were provided ad libitum access to water and standard chow (No. 2916, Harlan Teklad) or fasted for 24 hours. All animals in experiments were included. The ARRIVE guidelines 2.0 were used for reporting of the research (14).

### Immunoprecipitation of Tagged Mitochondria

Cell specific mitochondria were isolated as described in Bayraktar EC et. al. (10). All steps were performed at 4°C or on ice. Mouse renal tissue was homogenized with a Glas Col homogenizer (099C K54) at 4,000 RPM in either 1 mL of KPBS (136 mM KCl, 10 mM KH_2_PO_4_ pH 7.25) for metabolomics and immunoblot analyses or 1 mL mitochondrial isolation buffer (5 mM HEPES, 70 mM Sucrose, 220 mM Mannitol, 5 mM MgCl_2_, 10 mM KH_2_PO_4_, 1 mM EGTA, 1% fatty acid free BSA, pH 7.2) for respiration analyses. Homogenates were centrifuged for 5 minutes at 1000xg. 1 mL of supernatant was incubated with 30 μL of anti-HA beads (Thermo Scientific; 88837) for up to 10 minutes (3 minute minimum) with rotation at 4°C. HA-bead bound mitochondria were washed 2 times with 1 mL assay buffer, followed by assay specific dilution (described in the sections below).

### Isolation of Mitochondria from Whole Kidney

Mouse renal tissue was homogenized in 1 mL mitochondrial isolation buffer for respiration analysis (see above) or 1mL mitochondrial isolation buffer without BSA for immunoblot assays. Homogenates were centrifuged for 5 minutes at 600xg, followed by transfer of the supernatant into a new tube. This process was repeated 6 times. The transferred supernatant was then centrifuged for 10 minutes at 10,000xg, followed by removal of the supernatant. The remaining pellet was washed two times with 1 mL of mitochondrial isolation buffer followed by centrifugation at 10,000xg for 5 minutes at 4°C. After the second wash, the pellet was resuspended in assay specific buffer.

### Antibodies

Primary antibodies used include, Calnexin (Abcam Cat# ab22595, RRID:AB_2069006); CPT1A, (Proteintech Cat# 66039-1-Ig, RRID:AB_11041710); HDAC1 (Cell Signaling Technology Cat# 5356, RRID:AB_10612242); HSP90 (Bio-Rad; VMA00082); SDHA (Cell Signaling Technology Cat# 11998, RRID:AB_2750900); SGLT2 (Abcam Cat# ab85626, RRID:AB_10674183); NCC-DL488 (StressMarq Biosciences Cat# SPC-402D-DY594, RRID:AB_2825469); PMP70 (Abcam Cat# ab85550, RRID:AB_10672335); and VDAC (Cell Signaling Technology Cat# 4661, RRID:AB_10557420).

Secondary antibodies used include: donkey anti-rabbit AF488 (Thermo Fisher Scientific Cat# A-21206, RRID:AB_2535792); goat anti-chicken AF405 (Thermo Fisher Scientific Cat# A48260, RRID:AB_2890271); donkey anti-mouse AF488 (Thermo Fisher Scientific Cat# A-21202, RRID:AB_141607), donkey anti-rabbit AF594 (Thermo Fisher Scientific Cat# A-21207, RRID:AB_141637), Anti-mouse IgG, HRP-linked Antibody (Cell Signaling Technology Cat# 7076, RRID:AB_330924), and Anti-rabbit IgG, HRP-linked Antibody (Cell Signaling Technology Cat# 7074, RRID:AB_2099233).

### Immunoblotting

Mitochondria bound to anti-HA beads were incubated with triton lysis buffer (50 mM Tris HCl, 2% SDS, pH 6.8) supplemented with Halt Protease Inhibitor Cocktail (Thermo Scientific; 78443) for 20 minutes, followed by magnetization and removal of beads. Mitochondria extracted from whole tissue homogenates were resuspended in RIPA Lysis buffer (Teknova R3792) supplemented with Halt Protease Inhibitor Cocktail and incubated for 10 minutes. Protein concentration was evaluated using the DC Protein Assay kit (Bio-Rad; 5000112). Four micrograms of protein/sample were loaded into 4%– 20% Mini-PROTEAN TGX stain-free polyacrylamide gels (Bio-Rad; 4568093), transferred onto PVDF membranes (Bio-Rad; 1704273), blocked in 5% milk in 1x Tris-buffered saline with 1% Tween for 1 hour, and incubated with primary antibody overnight at 4°C. Membranes were then washed and then incubated with a secondary antibody for 1 hour at room temperature. Either fluorescent secondary antibodies or horseradish peroxidase linked secondary antibodies with chemiluminescence (Clarity Western ECL Substrate, Bio-Rad) were used. Blots were imaged with ChemiDoc MP (Bio-Rad). Densitometry determined based on total protein quantified using Image Lab (Bio-Rad).

### Immunostaining

Renal tissue was harvested and fresh frozen in Optimal Cutting Temperature (OCT) compound. 6 μm kidney sections were fixed with 4% paraformaldehyde for 30 minutes and simultaneously blocked and permeabilized in 5% normal donkey serum with 0.3% Triton-100 for 1 hour at room temperature. Tissue slides were incubated overnight with primary antibody in blocking solution overnight at 4°C, followed by incubation with secondary antibody, DAPI (Invitrogen 62248) and/or Lotus lectin flourescein (Vector Laboratories FL 13212) in blocking solution at 1:500, 1:5000, and 1:100 dilutions, respectively, for 1 hour at room temperature. Images of immunostained sections were acquired using the LSM 880 confocal microscope (Zeiss) or a Nikon Eclipse 80i microscope using a Nikon DS-Fi3 camera. Images were processed using ImageJ2 using the Fiji image processing package.

### Electron Microscopy

Mouse kidney tissue was processed using a modified Ellisman protocol, (https://dx.doi.org/10.17504/protocols.io.36wgq7je5vk5/v2). Briefly, thick 1 μm sections were cut using an RMC ultramicrotome, stained with toluidine blue for morphology evaluation, and thin 80 nm sections were imaged under a JEOL 1011 TEM at 80 KV. Instead of acetone, samples were embedded using ethanol. Kidney EM images were reviewed by a renal pathologist (BN).

### Mitochondrial Membrane Potential

Mitochondrial membrane potential was assessed in isolated mitochondria using the fluorescent dye tetramethylrhodamine ethyl ester (TMRE). Mitochondria were isolated from kidney tissues via immunoprecipitation of tagged mitochondria, as described above. An aliquot of the isolated mitochondria was used to determine protein concentration using the DC Protein Assay kit (Bio-Rad). A total of 5 μg of mitochondrial protein per well was plated in quadruplicate onto a black, clear-bottom 96-well plate, with two replicates treated with 5 μM carbonyl cyanide m-chlorophenylhydrazone (CCCP) for 10 min as controls to confirm loss of mitochondrial membrane potential. TMRE was added to a final concentration of 100 nM, and samples were incubated at 37°C for 20 minutes in the dark. Following incubation, fluorescence was measured using a microplate reader (excitation/emission: 549/575 nm).

### Mitochondrial Respiration

Mitochondrial respiration was measured using the Agilent Seahorse (Seahorse XFe96 Extracellular Flux Analyzer) via the UT Southwestern Children’s Medical Center Research Institute Metabolomics Core. An aliquot of isolated mitochondria either bound to HA-beads or extracted from whole tissues homogenates were evaluated for protein concentration using the DC Protein Assay kit (Bio-Rad). Specific amounts of mitochondria based on assayed substrates were then resuspended in 20 μL of mitochondrial isolation buffer and plated onto a 96-well plate: 2 μg/well for Complex I activity (10 mM pyruvate/1 mM malate); 1 μg/well for Complex II function (5 mM succinate/2 μM rotenone); 2 μg/well for CPT1-independent FAO (50 μM palmitoylcarnitine/1 mM malate); 2.5 μg/well for CPT1-dependent FAO (40 μM Palmitoyl-CoA/0.5 mM carnitine/1 mM malate). After plating mitochondria, the 96-well plate with added mitochondria was centrifuged for 2 minutes at 2,700xg at 4°C. Buffer with or without substrate mixture (130 μL) was added to each well using 4 technical replicates per sample. Oxygen consumption rate (OCR) was measured as follows: baseline measurements were taken for 6 minutes; followed by treatment with 4 mM adenosine diphosphate (ADP) and measurement for 3 minutes; followed by treatment with 2 μM Oligomycin and measurement for 5 minutes; followed by treatment with 2 μM CCCP and measurement for 3 minutes; followed by treatment with either 2 μM Antimycin A or 10 μM Cytochrome C. Subsequent analysis was performed using the Agilent Wave XF Software v2.6.1. OCR of samples without substrate mixture (isolation buffer only) was considered background activity and was subtracted from OCR of respective samples with substrate. Data shown as individual biological replicates and representative of at least 2 independent experiments.

### Metabolomics

HA-bead bound mitochondria were incubated in LC/MS extraction buffer (80% methanol/20% LC/MS water) and samples frozen at -80°C until targeted screening metabolomic analysis was performed by the UT Southwestern Children’s Medical Center Research Institute Metabolomics Core. Data acquisition was performed by reverse-phase chromatography on a 1290 UHPLC liquid chromatography (LC) system interfaced to a high-resolution mass spectrometry (HRMS) 6550 iFunnel Q-TOF mass spectrometer (MS) (Agilent Technologies, CA). The MS was operated in both positive and negative (ESI+ and ESI-) modes. Analytes were separated on an Acquity UPLC® HSS T3 column (1.8 μm, 2.1 x 150 mm, Waters, MA). The column was kept at room temperature. Mobile phase A composition was 0.1% formic acid in water and mobile phase B composition was 0.1% formic acid in 100% ACN. The LC gradient was 0 min: 1% B; 5 min: 5% B; 15 min: 99%; 23 min: 99%; 24 min: 1%; 25 min: 1%. The flow rate was 250 μL min-1. The sample injection volume was 5 μL.

ESI source conditions were set as follows: dry gas temperature 225 °C and flow 18 L min-1, fragmentor voltage 175 V, sheath gas temperature 350 °C and flow 12 L min-1, nozzle voltage 500 V, and capillary voltage +3500 V in positive mode and −3500 V in negative. The instrument was set to acquire over the full m/z range of 40–1700 in both modes, with the MS acquisition rate of 1 spectrum s-1 in profile format.

Raw data files (.d) were processed using Profinder B.08.00 SP3 software (Agilent Technologies, CA) with an in-house database containing retention time and accurate mass information on 600 standards from Mass Spectrometry Metabolite Library (IROA Technologies, MA) which was created under the same analysis conditions. The in-house database matching parameters were: mass tolerance 10 ppm; retention time tolerance 0.5 min. Peak integration result was manually curated in Profinder for improved consistency and exported as a spreadsheet (.csv). Data normalized by the total ion count (**Supplemental Table 1**). Metabolite isoforms with identical molecular weights were summed and treated as a single species, and a square root transformation was subsequently applied for all downstream analysis. Single and two variable principal component analysis (PCA) plots were generated using Metabolanalys.ca. Total acylcarnitines per sample were calculated as the sum of all acylcarnitine species detected (Acetylcarnitine, Carnitine (6:0), Carnitine (8:0), Decanoylcarnitine, Carnitine (12:0), trans-2-Dodecenoylcarnitine, trans-2-Tetradecenoylcarnitine, Carnitine (16:0), Stearoylcarnitine, and Carnitine (18:1)).

### Plasma creatinine measurements

Retroorbital or submandibular blood was harvested and plasma was isolated using lithium heparin-coated plasma separator tubes (BD). Plasma creatinine was assayed using liquid chromatography-mass spectrometry by the O’Brien Center for Acute Kidney Injury Research, Pre-clinical Studies of AKI Biomedical Resource Core at the University of Alabama at Birmingham.

### RNA extraction and quantification

For RNA extraction, kidneys were harvested into RNA Bee isolation reagent and disrupted by bead homogenization in Fisherbrand Bead Mill Tubes using a Fisherbrand Bead Mill 24 Homogenizer. RNA was extracted using Direct-Zol (Zymo Research) kits according to the manufacturer’s protocol. cDNA synthesis was performed using MMLV reverse transcriptase (Clontech) with oligo(dT) primers. qRT-PCR reactions were performed on a QuantStudio7 Flex (Applied Biosystems) using iTaq^TM^ Universal SYBR Green Supermix (Bio-Rad). *Cpt1a* transcript levels were normalized to *Rpl13a*. Primers used include: *Cpt1a* Fw: TTATGTGAGTGACTGGTGGGAGG Rev: ATGGGTTGGGGTGATGTAGAGC, *Rpl13a* Fw: GAGGTCGGGTGGAAGTACCA Rev: TGCATCTTGGCCTTTTCCTT.

### Statistical Analysis

Statistical analyses were performed using Prism 10.0 (GraphPad). One-way ANOVA with Dunnett’s multiple comparison analysis and Two-way ANOVA with Sidak’s multiple comparison analysis was used to compare mitochondrial OCR and western blot protein densitometry across groups and conditions. Statistical analysis for metabolomics described above. A *P* value < 0.05 was considered statistically significant. Prism 10.0 (GraphPad), Illustrator (Adobe), PowerPoint (Microsoft), and BioRender.com were used to generate figures.

## Results and Discussion

### Isolation of kidney cell-specific mitochondria from cortical tubular cells

The MITO-Tag mouse model (10) was developed to isolate mitochondria from tissues in a cell-specific manner. Cre recombinase activity leads to expression of an HA-tagged construct of a mitochondrial outer membrane protein (3xHA-eGFP-OMP25) allowing for immuno-isolation of mitochondria using anti-HA beads (**Figure 1C**). In comparison to whole tissue mitochondrial isolation which requires multiple centrifugation steps and could require up to an hour, MITO-Tag mitochondria isolation could be accomplished within minutes. In order to tag mitochondria from the early PT, we generated the *Slc34a1-CreERT2;MITO-Tag* (Slc34a1-MT) mouse model. The *Slc34a1-CreERT* inducible model (11) with tamoxifen dose dependent efficiency allows for targeting the early S1-S2 segment of the kidney with one dose of tamoxifen. In contrast to the MitoTag (Rosa26-CAG-LSL-GFP-OMM) mouse (15), the EGFP of the MITO-Tag (3xHA-eGFP-OMP25) mouse does not fluoresce. However, it is possible to immunostain tissue using anti-GFP antibodies. Slc34a1-MT kidneys showed co-localization of MITO-Tag expression in predominantly PT cells that express sodium glucose co-transporter 2 (SGLT2) in the early S1-S2 segment (**Figure 1D**). The MITO-Tag in Slc34a1-MT kidneys also co-localized to lotus tetragonolobus lectin (LTL), a pan PT stain, but did not co-localize with the DCT-specific transporter, the sodium-chloride co-transporter (NCC). The *Ggt1-Cre;MITO-Tag* (Ggt1-MT) mouse model (12) was used to tag mitochondria from the late PT, the S2-S3 segment (**Figure 1D**). In the Ggt1-MT kidneys, MITO-Tag co-localized with LTL, but did not co-localize with SGLT2-expressing cells, supporting the expression of MITO-Tag in the late PT. We next generated the *PvAlb-Cre;MITO-Tag* (PvAlb-MT) mouse model which tags early DCT mitochondria (16) (**Figure 1D**). In PvAlb-MT kidneys, MITO-Tag expression co-localized with the DCT-specific NCC, but not with any of the PT markers.

Addition of transgene tags has the potential of causing cellular dysfunction. In fact, introduction of the MitoTag (Rosa26-CAG-LSL-GFP-OMM) transgene in B6.Cg-Tg(NPHS2-cre)295Lbh/J mice to tag podocyte mitochondria was reported to result in podocyte injury and abnormal mitochondrial aggregation leading to renal failure and premature mortality (17). In contrast, expression of MITO-Tag (3xHA-eGFP-OMP25) in podocytes did not result in podocyte injury or mitochondrial abnormalities (17). Thus, we next assessed whether the addition of the MITO-Tag (3xHA-eGFP-OMP25) protein on the outer membrane of the mitochondria would affect mitochondrial structure and kidney function when expressed in the PT and DCT. Transmission electron microscopy showed that in all three models, the addition of MITO-Tag in the respective tubular segments did not alter mitochondrial morphology (**Figure 2A**). Baseline renal function as measured by plasma creatinine were within normal reference range (18–21) (**Table 1**), suggesting that expression of MITO-Tag (3xHA-eGFP-OMP25) in the PT or DCT did not disrupt mitochondrial morphology or renal function. Using anti-HA immunoprecipitation, all three models showed enrichment of mitochondria, but not nuclear or cytoplasmic components (**Figure 2B**). As contact sites are well known between mitochondria and other organelles, particularly the endoplasmic reticulum (ER) and peroxisomes (22), we next assessed the extent to which the anti-HA immunoprecipitation co-isolates mitochondria with other organelles. Similar to the hepatic MITO-Tag model (10) and traditional whole tissue mitochondrial isolation, anti-HA MITO-Tag immunoprecipitation with our renal MITO-Tag models also enriched for both ER and peroxisomes (**Figure 2C**). Immunostaining showed co-localization of MITO-Tag expression with calnexin (ER) and PMP70 (peroxisome) in Slc34a1-MT, Ggt1-MT, and PvAlb-MT kidneys (**Figure 2D**).

**Figure 2.**
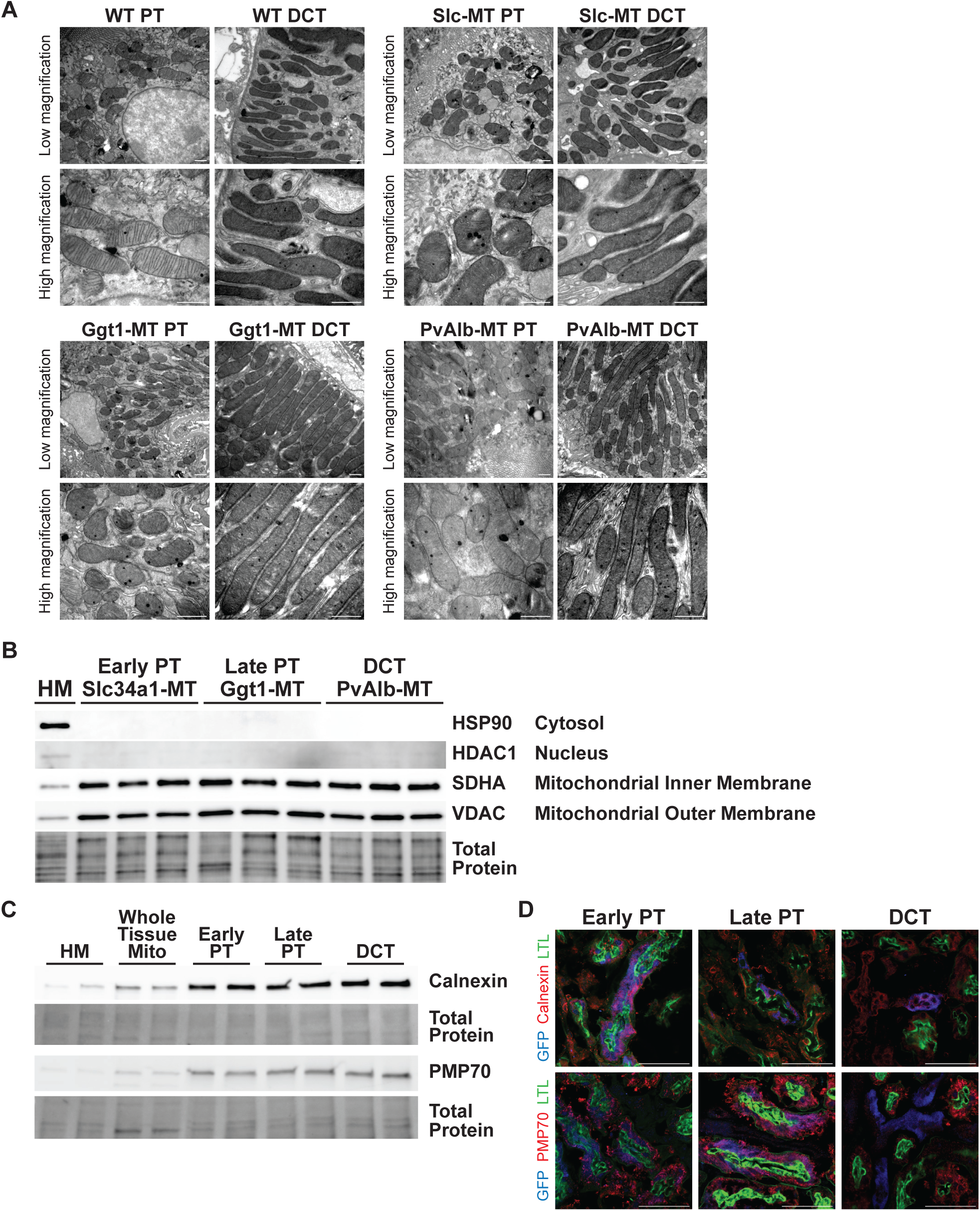
Isolation of intact kidney cell specific mitochondria. **A.** Electron microscopy of proximal tubule and distal convoluted tubules at low and high magnification of wild-type (WT, Cre negative), Slc34a1-MT, Ggt1-MT, and PvAlb-MT kidneys. Low magnification, 20000X. High magnification, 50000X. Scale bar = 1 μm. **B.** Immunoblot of protein samples of whole kidney homogenate (HM) and mitochondria isolated via anti-HA immunoprecipitation from Slc34a1-MT, Ggt1-MT, and PvAlb-MT kidneys. **C.** Immunoblot of protein samples of whole kidney homogenate (HM), mitochondria isolated from whole kidney (whole tissue mito), and mitochondria isolated via anti-HA immunoprecipitation from Slc34a1-MT, Ggt1-MT, and PvAlb-MT kidneys. **D.** Immunostaining with anti-GFP (MITO-Tag), Calnexin (endoplasmic reticulum), PMP70 (peroxisome), and LTL of Slc34a1-MT, Ggt1-MT, and PvAlb-MT kidneys. Scale bar = 50 μm.

**Table 1.** Mouse model plasma creatinine.

| Mouse line | Plasma Creatinine (mg/dL) |
| --- | --- |
| Early PT (Slc34a1-MT) mice | 0.075 ± 0.005 |
| Late PT (Ggt1-MT) mice | 0.093 ± 0.022 |
| DCT (PvAlb-MT) mice | 0.095 ± 0.027 |

DCT, distal convoluted tubule; MT, MITO-Tag; PT, proximal tubule.
Data shown as mean ± SD.

### Differential mitochondrial functional capacities

In order to test mitochondrial viability, we compared the respiratory response to cytochrome c between MITO-Tag isolated mitochondria and whole kidney mitochondria isolated using traditional methods. Similar to whole kidney isolated mitochondria, the MITO-Tag isolated mitochondria exhibited a minimal increase in respiration in the presence of cytochrome c (23), suggesting that the mitochondrial outer membrane using the rapid MITO-Tag isolation protocol remained largely intact (**Figure 3A**). We next assessed the mitochondrial membrane potential of the MITO-Tag isolated kidney mitochondria. Interestingly, the mitochondrial membrane potential was higher in the DCT compared to that of the proximal tubule (**Figure 3B**). This difference in mitochondrial membrane potential is in line with prior reports showing higher mitochondrial membrane potential in the distal tubule (thick ascending limb) compared to that of the proximal tubule (7), suggesting that mitochondrial membrane potential varies in the different tubular cells.

**Figure 3.**
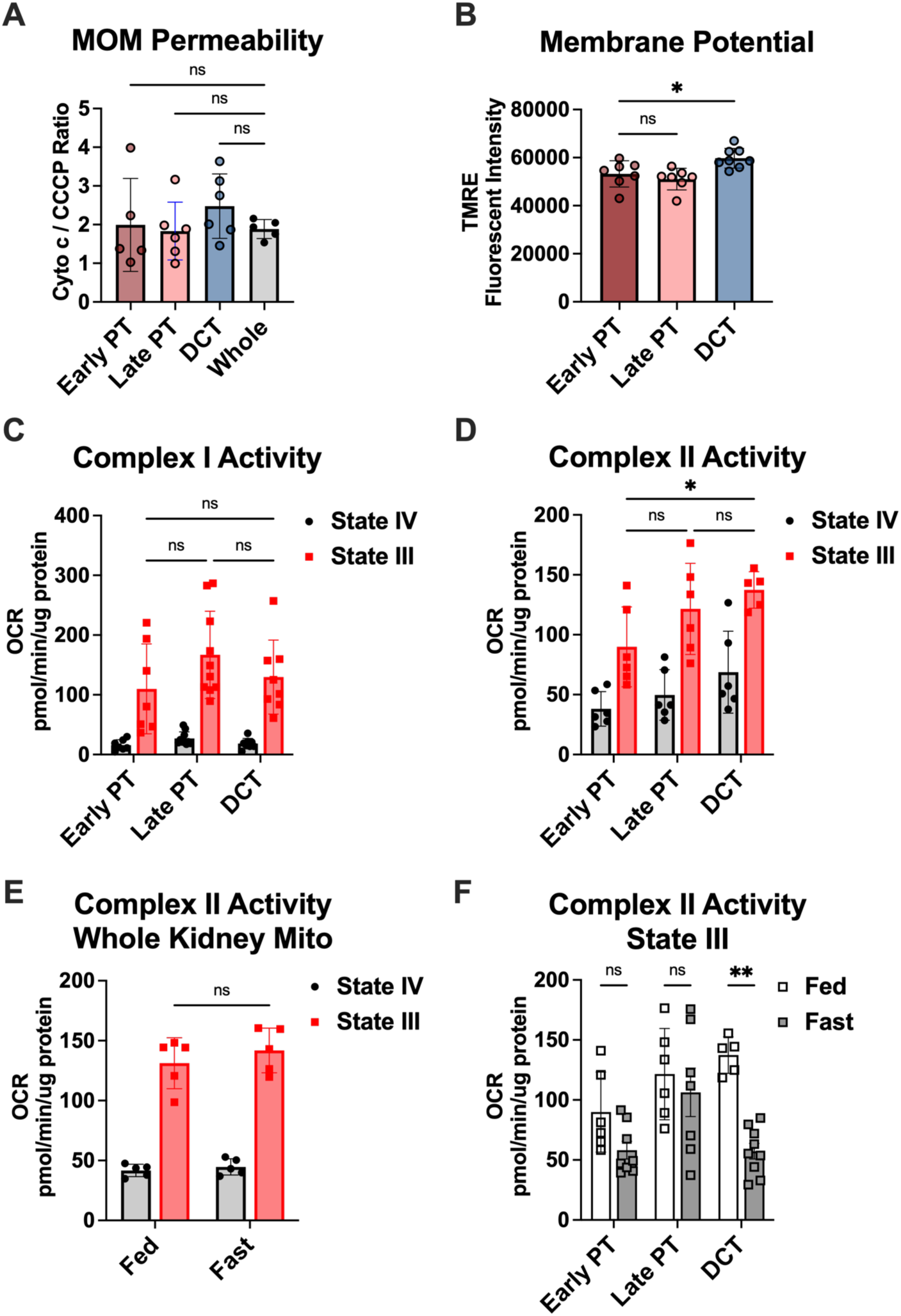
MITO-Tag isolated kidney cell-specific mitochondria are viable and respirating. Kidney mitochondria were isolated via traditional methods (whole) or via anti-HA immunoprecipitation from Slc34a1-MT (Early PT), Ggt1-MT (Late PT), and PvAlb-MT (DCT) kidneys. Data shown as mean±SD. **A.** Mitochondrial outer membrane (MOM) permeability measured by oxygen consumption rate (OCR) using Complex II substrate (5 mM Succinate) with cytochrome c treatment after treatment with an uncoupler (CCCP). Data shown as ratio of cytochrome c OCR versus CCCP OCR. n=5-6/group. **B.** Mitochondrial membrane potential measured by TMRE fluorescent intensity. n=7-8/group. **C-F.** Mitochondrial respiration measured by oxygen consumption rate (OCR) before (State IV) and after 4 mM ADP (State III). **C.** Complex I activity of Slc34a1-MT (Early PT), Ggt1-MT (Late PT), and PvAlb-MT (DCT) mitochondria isolated from fed mice. n=7-10/group. **D.** Complex II activity of Slc34a1-MT (Early PT), Ggt1-MT (Late PT), and PvAlb-MT (DCT) mitochondria isolated from fed mice. n=5-6/group. **E.** Complex II activity of whole kidney mitochondria isolated from fed or fasted mice. n=5/group. **F.** Complex II activity of Slc34a1-MT (Early PT), Ggt1-MT (Late PT), and PvAlb-MT (DCT) mitochondria isolated from fed or fasted mice. n=5-9/group. **A,B.** One-way ANOVA with Dunnett’s multiple comparisons test, ns, not significant. Bartlett’s test for homogeneity p = 0.0729. **C-F.** Two-way ANOVA with Sidak’s multiple comparisons test, \**P*<0.05, \*\**P*<0.01.

As mitochondrial function has been described to be unique across organs (1), we next examined whether the functional capacities of mitochondria also vary across different tubular segments. To assess mitochondrial function, we isolated Slc34a1-MT, Ggt1-MT, and PvAlb-MT mitochondria and measured their respiratory capacity, comparing oxygen consumption without ADP (State IV respiration) and with ADP (State III respiration). We observed robust induction of mitochondrial respiration with addition of ADP when measuring Complex I and II activity (**Figure 3C,D**). Across the three renal tubular cortical segments, there were no significant differences in Complex I activity, while the early PT exhibited the lowest Complex II activity. As it has been described that liver mitochondria decrease Complex II activity after an overnight fast (24), we next measured kidney mitochondrial respiratory capacities after 24 hours of fasting. At the whole kidney level, isolated bulk kidney mitochondria exhibited no significant difference in Complex II activity between fed and fasted states (**Figure 3E**). However, when comparing mitochondrial Complex II activity across the different tubular segments, fasting did not change Complex II activity in late PT mitochondria, however, similar to liver mitochondria, fasting significantly decreased Complex II activity in DCT mitochondria and trended lower in early PT mitochondria (**Figure 3F**).

Given the dynamic changes in mitochondrial respiratory capacity after a metabolic stress, we next performed metabolomic analysis on segment-specific MITO-Tag isolated mitochondria in fed and fasted conditions. Untargeted screening metabolomics identified 125 metabolites (**Supplemental Table 1**). By principal component analysis (PCA), global metabolite signatures were similar across the early PT, late PT, and DCT mitochondria when analyzed within fed or fasted states (**Figure 4A**). Two factor PCA analysis suggests that early PT, late PT, and DCT mitochondria respond similarly to fasting (**Figure 4B**). As fasting is known to alter acylcarnitines levels in a tissue-specific manner (25), we next analyzed acylcarnitines specifically in the kidney cell specific mitochondria. While some shorter chain acylcarnitines were increased during fasting across all three segments, other medium and longer chain acylcarnitines trended higher in response to fasting in the late PT and DCT mitochondria while they remained unchanged in the early PT (**Figure 4C**). In the late PT and DCT, total acylcarnitine mitochondrial content increased, while free carnitine was significantly decreased during fasting. In contrast, the early PT mitochondria exhibited a variable non-significant trend in total acylcarnitine and free carnitine abundances.

**Figure 4.**
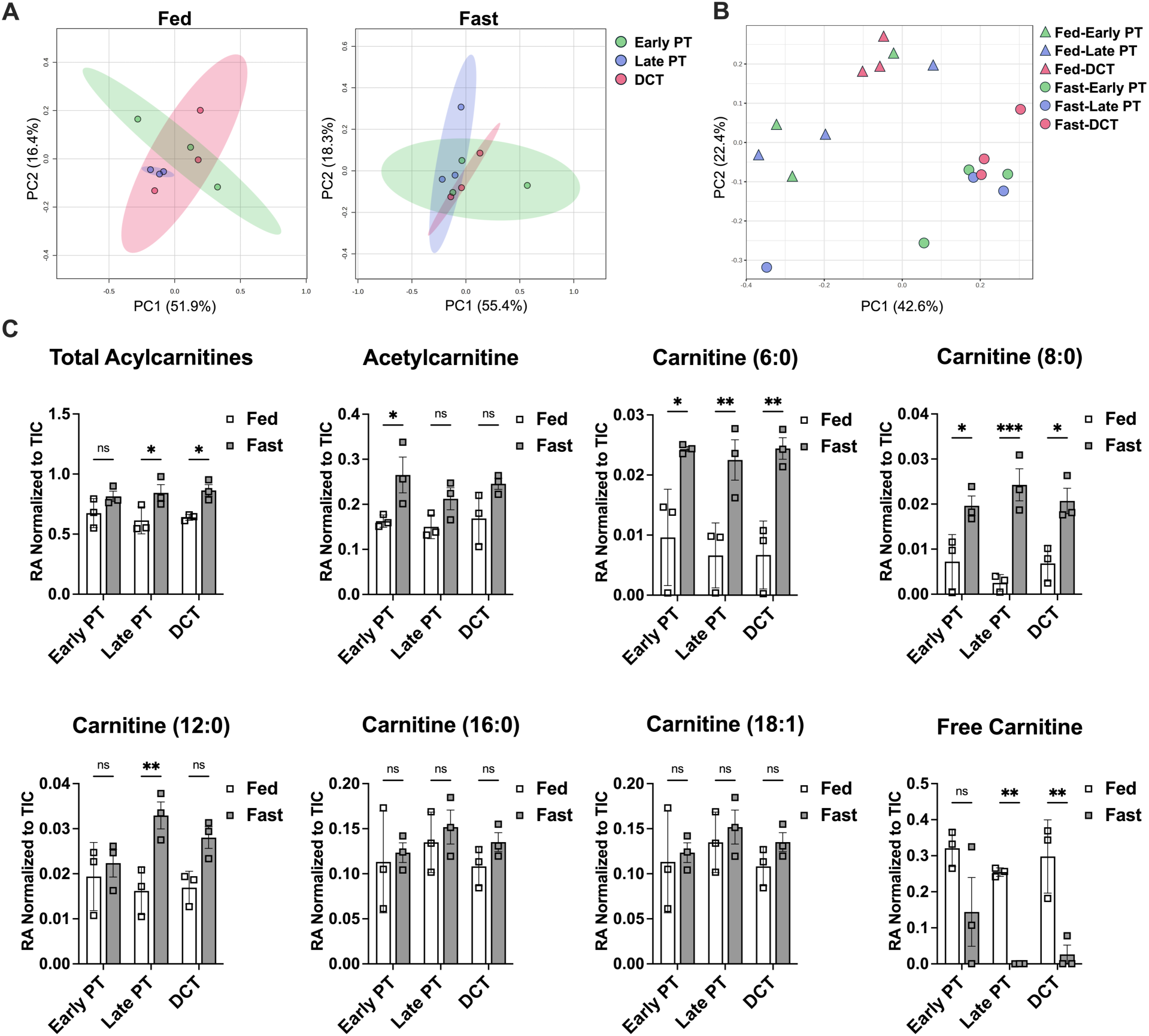
Differential and fasting-induced changes in mitochondrial metabolite composition in the proximal tubule and the distal convoluted tubule. Mitochondria isolated via anti-HA immunoprecipitation from Slc34a1-MT (Early PT), Ggt1-MT (Late PT), and PvAlb-MT (DCT) kidneys from fed or 24-hour fasted mice were submitted for targeted screening metabolomics by LC/MS-MS, n=3/group. Analysis performed via Metaboanalyst, details described in Methods. **A.** Principal components analysis (PCA) of metabolites across the early PT, late PT, and DCT in fed and fasted conditions. **B.** Two factor PCA of metabolites across the early PT, late PT, and DCT in fed and fasted conditions. **C.** Relative abundance (RA) of total acylcarnitines, individual acylcarnitines and acetylcarnitine, and free carnitine. Data shown normalized to total ion count (TIC). Data shown as mean±SD. Two-way ANOVA with Sidak’s multiple comparisons test, \**P*<0.05, \*\**P*<0.01, \*\*\**P*<0.001, ns, not significant.

### Measuring renal cell-specific fatty acid oxidation

As acylcarnitines are key starting substrates for FAO (26), we next examined FAO capacity in the late PT and DCT given the notable increase in total acylcarnitines and decrease in free carnitine during fasting. Using palmitoyl-CoA as an FAO substrate (**Figure 5A**), we compared the difference in palmitoyl-CoA oxygen consumption of mitochondria isolated from a whole kidney homogenate using traditional methods and immunoprecipitated late PT- and DCT-specific mitochondria in response to fasting, a state that will increase FAO (27). Similar to bulk mitochondria isolated from the whole kidney, DCT-specific mitochondria exhibited a modest non-significant increase in palmitoyl-CoA oxidation in the fasted state compared to the fed state (**Figure 5B**). In contrast, late PT mitochondria, exhibited a robust increase in palmitoyl-CoA induced oxygen consumption.

**Figure 5.**
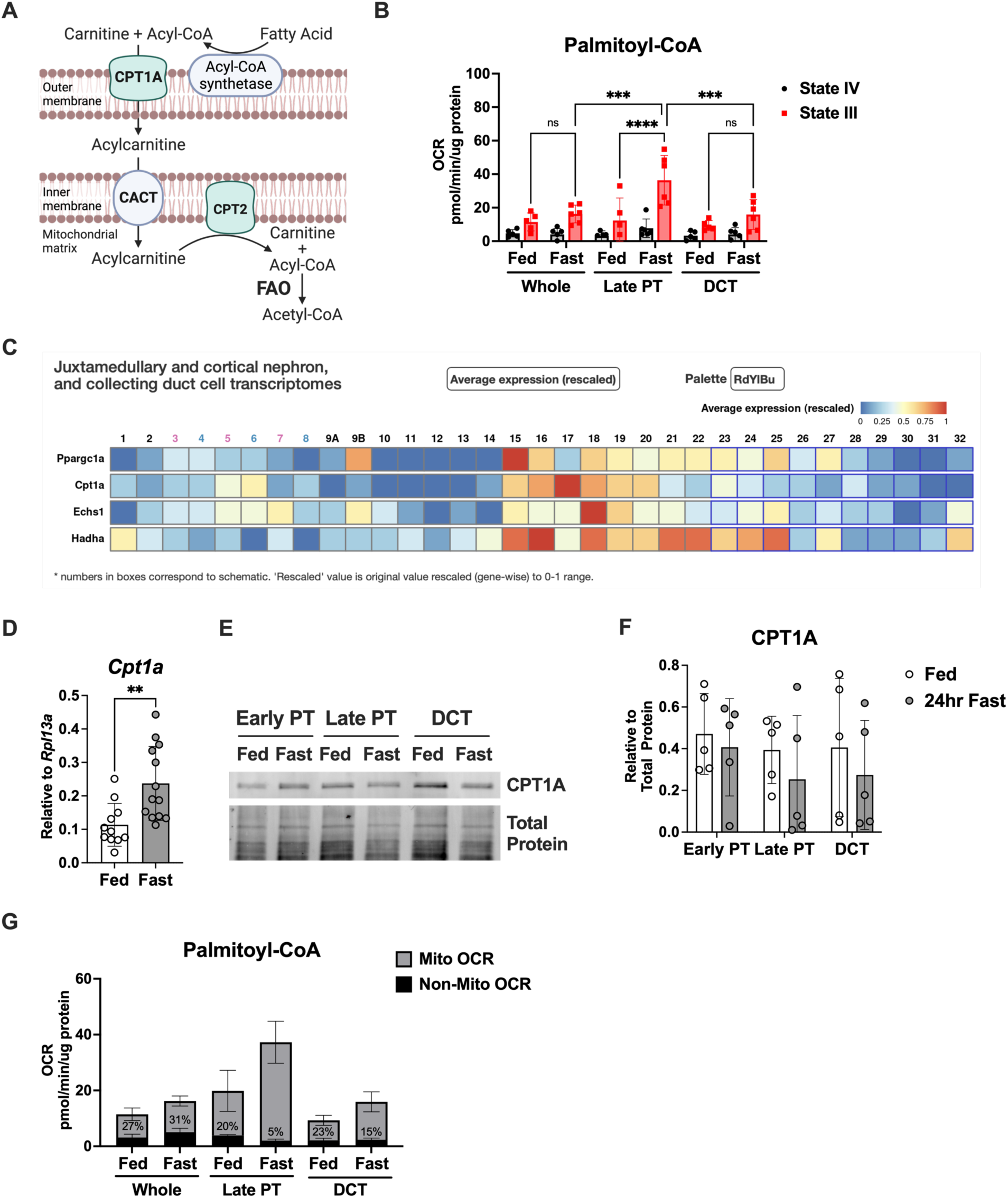
Differential fatty acid oxidation capacities in late proximal tubular and distal convoluted tubular mitochondria in response to fasting. **A.** Diagram of fatty acid oxidation. Created in BioRender. Huen, S. (2025) https://BioRender.com/9rvibxd **B.** Mitochondria were isolated via traditional whole kidney tissue isolation (whole) or anti-HA immunoprecipitation from Ggt1-MT (Late PT) and PvAlb-MT (DCT) kidneys from fed or 24-hour fasted mice. FAO capacity measured by OCR using 40 μM palmitoyl-CoA as a substrate, before (State IV) and after 4 mM ADP (State III), n=5-6/group. **C.** Single cell RNA-sequencing data from Ransick et al. (5) of select FAO genes visualized via Kidney Cell Explorer at https://cello.shinyapps.io/kidneycellexplorer/. 3/4: S1 PT female/male, 5/6: S2 PT female/male, 7/8: S3 PT female/male, 18: DCT. **D.** Whole kidney mRNA expression shown relative to *Rpl13a*. n=11-13/group. **E.** Representative immunoblot of protein samples of mitochondria isolated via anti-HA immunoprecipitation. **F.** Densitometry of (E), n=5/group, no comparison was significantly different. **G.** After ADP-stimulation, OCR measured after treatment with antimycin A, which specifically inhibits mitochondrial function, is the non-mitochondrial contribution to OCR. Percentages shown on bar graph represent the percent contribution of non-mitochondrial OCR. Related to Figure 5B. **B,F.** Data shown as mean±SD. Two-way ANOVA with Sidak’s multiple comparisons test, \*\*\**P*<0.001, \*\*\*\**P*<0.0001. **D.** Data shown as mean±SD. Unpaired t-test, \*\**P*<0.01.

Classic microdissection studies of tubular segments measuring ATP production in response to single substrates found that the PT preferentially uses fatty acids for energy as compared to the DCT which uses glucose, lactate and ketones (6). However, the DCT expresses genes involved in FAO (6, 28, 29). Carnitine palmitoyltransferase 1 (CPT1) is often considered a rate-limiting step for FAO. Long chain acyl-CoAs such as palmitoyl-CoA require CPT1 activity to enter the mitochondria for beta oxidation (**Figure 5A**). The predominate form of CPT1 in the kidney is CPT1A. At baseline, single cell RNA-Sequencing data show that mitochondrial FAO related genes including *Cpt1a* are more abundant in the DCT than in any segment of the PT ((5) and **Figure 5C**). At a whole kidney level, *Cpt1a* transcript increased in response to fasting (**Figure 5D**). However, at the level of cell specific mitochondria, CPT1A protein expression was not significantly different across the early PT, late PT, and DCT, and did not change in response to fasting. These data suggest that a change in FAO capacity involving CPT1A activity can be independent of protein abundance (**Figure 5E,F**). These data highlight the power of this kidney cell-specific mitochondrial system, as differences in FAO rates and capacity at a kidney cell-specific level and the magnitude of the changes that occur in response to a metabolic stress cannot be captured by a mixed population of isolated mitochondria at the whole kidney level and may not be reflective of gene transcript or protein expression.

## Discussion

The application of the MITO-Tag system to kidney tubular epithelial cells provides the opportunity to interrogate mitochondrial metabolism in a cell-specific manner. Here we present novel data suggesting that in response to the acute metabolic stress of fasting, FAO capacity differentially changes between the late PT mitochondria compared to DCT mitochondria. The advantage of this system allows for cell specificity and focuses the metabolic analyses to mitochondria, organelles that are not only the power generators for cellular metabolism, but also the site where the rate-limiting steps of fuel utilization, ketogenesis, and gluconeogenesis occur. Moreover, the renal MITO-Tag system can measure the functional capacities of cell-specific mitochondria and elucidate metabolic properties that may not be reflected by gene transcript or protein abundances.

While the kidney MITO-Tag system can elucidate the mitochondrial heterogeneity across the diverse landscape of the many kidney cell types, there are some limitations to our model. First, co-isolation with other organelles can be a limitation, however, this is inherent to the biology of contact sites known to exist and associate mitochondria with other organelles. As peroxisomes are co-isolated with the MITO-Tag system, the metabolomics and FAO measurements cannot fully differentiate between mitochondrial and peroxisomal metabolism. Moreover, peroxisomes are more abundant in the PT (**Figure 2D**) and could compensate in the context of tubular CPT1A deficiency (30). While it has been described that peroxisomes can accept palmitoylcarnitine as a substrate in the context of mitochondrial FAO defects, peroxisomal oxidation of palmitoylcarnitine is estimated at only ∼2% of whole cell activity (31). Moreover, the level of non-mitochondrial oxygen consumption in response to palmitoyl-CoA was similar, if not lower, in the MITO-Tag isolated mitochondria compared to that of mitochondria isolated from the whole kidney using traditional methods (**Figure 5G**). Interestingly, the contribution of non-mitochondrial oxygen consumption in MITO-Tag isolated mitochondria from fasted mice is even lower, suggesting that the increase in FAO capacity observed in late PT in response to fasting is driven primarily from mitochondria. Thus, our findings suggesting differential acyl-CoA FAO metabolism in the context of fasting is likely driven primarily by mitochondrial metabolism, with a potential minor contribution from peroxisomes. Second, we leveraged the tamoxifen dose dependence to enrich for early S1-S2 PT cells, however, the *Slc34a1-CreERT2* and *Ggt1-Cre* are not exact in their targeting of the S1, S2, and S3 segments of the proximal tubule. Additionally, tamoxifen and the oil vehicle could have effects on mitochondrial metabolism. The *PvAlb-Cre* driver only targets the early DCT. Future work is needed to investigate the metabolic differences between DCT1 and DCT2, as well as other renal cell types. Third, the studies presented here were performed in male mice. Future work will investigate kidney cell-specific sexual dimorphism of mitochondrial metabolism which has been reported in other tissues (32, 33). Finally, inherent to the MITO-Tag method is the removal of the mitochondria from the in vivo environment. As mitochondria are isolated from the cell and are the focus of this approach, this system cannot examine whole cell metabolism such as glycolysis. Future research is needed to incorporate technologies such as MALDI-MSI (34) to integrate spatial metabolomics with ex vivo renal MITO-Tag functional studies.

## Conclusion

The renal MITO-Tag model has the power of interrogating kidney cell-specific mitochondrial function. This ability to measure kidney cell-specific metabolism can be transformative and enable the advancement in several key areas of kidney research, including kidney development, physiology, disease pathogenesis and mitochondrial biology and therapeutics. Using this system, we have for the first time defined the mitochondrial metabolic heterogeneity of renal cortical tubular epithelium and discovered differential FAO capacities of late PT and the DCT.

## Data Statement

All data generated during this study are included in this article or in the Supplemental Materials. Metabolomic data are included in Supplemental Table 1.

## Acknowledgements

The authors wish to thank Dr. Mariya Sweetwyne for helpful discussions. Preprint is available at https://doi.org/10.1101/2024.11.24.622516.

## Author Contributions

Conceptualization/Design: K.F., A.H.V., S.C.H., Investigation: K.F., A.H.V., M.R., J.D., T.B., C.L., D.S., Formal Analysis: K.F., A.V., M.R., P.M., B.N., S.C.H., Writing–Original Draft: S.C.H., Writing– Review/Editing: K.F., A.V., P.M., S.C.H. All authors approved the final manuscript.

---

**Supplemental Table 1. Targeted metabolomics of renal cortical cell-specific mitochondria in fed and fasted states.**

Mitochondria isolated via anti-HA immunoprecipitation from Slc34a1-MT (Slcfed, Slcfast), Ggt1-MT (Ggtfed, Ggtfast), and PvAlb-MT (PvAfed, PvAfast) kidneys from fed or 24-hour fasted mice were submitted for targeted screening metabolomics by LC/MS-MS, n=3/group. Data shown as normalized to total ion count.

